# Characterizing microRNA-mediated modulation of gene expression noise and its effect on synthetic gene circuits

**DOI:** 10.1101/2020.07.08.193094

**Authors:** Lei Wei, Shuailin Li, Tao Hu, Michael Q. Zhang, Zhen Xie, Xiaowo Wang

## Abstract

Gene expression noise plays an important role in many biological processes, such as cell differentiation and reprogramming. It can also dramatically influence the behavior of synthetic gene circuits. MicroRNAs (miRNAs) have been shown to reduce the noise of lowly expressed genes and increase the noise of highly expressed genes, but less is known about how miRNAs with different properties may regulate gene expression noise differently. Here, by quantifying gene expression noise using mathematical modeling and experimental measurements, we showed that competing RNAs and the composition of miRNA response elements (MREs) play important roles in modulating gene expression noise. We found that genes targeted by miRNAs with weak competing RNAs show lower noise than those targeted by miRNAs with strong competing RNAs. In addition, in comparison with a single MRE, repetitive MREs targeted by the same miRNA suppress the noise of lowly expressed genes but increase the noise of highly expressed genes. Additionally, MREs composed of different miRNA targets could cause similar repression levels but lower noise compared with repetitive MREs. We further observed the influence of miRNA-mediated noise modulation in synthetic gene circuits which could be applied to classify cell types using miRNAs as sensors. We found that miRNA sensors that introduce higher noise could lead to better classification performance. Our results provide a systematic and quantitative understanding of the function of miRNAs in controlling gene expression noise and how we can utilize miRNAs to modulate the behavior of synthetic gene circuits.

## Introduction

Stochastic fluctuations lead to variation, or noise, in gene expression levels, which is inevitable even among genetically identical cells exposed to the same environmental conditions (1). In clonal populations of microbes, gene expression noise enables cells to generate diverse phenotypes, which may improve the fitness of the population in certain environments (2–5). In multicellular organisms, noise is involved in many biological processes, such as cell differentiation (6, 7), reprogramming (8), and apoptosis (9). Noise can give rise to the heterogeneity of cancer cells within individual tumors (10, 11). In addition, gene expression noise can influence the behavior of synthetic gene circuits. In transcriptional cascades, noise is amplified near the transition region of the dose-response curve, reducing the precision of signal transduction (12–15). Nonetheless, noise is necessary in excitable circuits for the initiation of transient state switching (3, 4, 16). Therefore, it is valuable to understand the mechanism of noise modulation in nature and thus to manipulate gene expression noise to control phenotypes according to expectations.

MiRNAs are ∼22 nt noncoding RNAs that mediate the posttranscriptional regulation of their target genes by either translational repression or poly(A)-tail shortening (17, 18). Recently, researchers proposed that miRNAs can control gene expression noise and confer robustness to biological processes (19–23). MiRNAs can reduce the noise of lowly expressed genes by accelerating mRNA turnover, which can be compensated for by higher transcription rates, and miRNAs can increase the noise of highly expressed genes by introducing additional extrinsic noise (20). These discoveries provided an explanation for the observation that many highly expressed housekeeping genes do not harbor miRNA binding sites (24). Furthermore, the capacity of miRNAs to modulate noise was found to be highly related to the strength of their repression on the expression of target genes (20).

MiRNAs are important regulators in both natural gene networks and synthetic gene circuits. Different types of miRNAs and various miRNA targets are involved in diverse biological processes. Therefore, it is essential to delineate how the properties of miRNAs and their targets beyond repression strength, such as the competing RNAs of miRNAs and the composition of miRNA binding sites, could modulate gene expression noise. For example, the competing RNAs of miRNAs have been shown to have the capacity to modulate gene expression noise (25, 26). Previously, we analyzed gene expression noise at the mRNA level by single-cell RNA-seq and revealed that miRNAs with weakly-interacted competing RNAs could buffer gene expression noise compared to miRNAs with few targets (27). However, there is a lack of comprehensive studies to systematically and quantitatively depict how the properties of miRNAs and the compositions of miRNA response elements (MREs) modulate gene expression noise and how such noise modulation further influences the output behavior of synthetic gene circuits. The absence of investigation hinders the understanding and utilization of miRNAs to control gene expression noise in natural gene regulatory networks and synthetic gene circuits.

Here, by quantifying gene expression noise via mathematical modeling and flow cytometry analysis with a dual-fluorescent reporter system, we investigated how competing RNA networks can influence gene expression noise at the protein level. Specifically, in comparison with genes targeted by miRNAs with strong competing RNAs, those targeted by miRNAs with weak competing RNAs show reduced gene expression noise over a wide range of gene expression levels. Furthermore, genes with repetitive MREs for the same miRNA exhibit lower noise than genes with a single MRE when the gene expression level is low but show a strong increase in noise when the gene expression level is high due to the saturation effect. Additionally, in comparison with repetitive MREs, MREs composed of targets that are regulated by different miRNAs exert a similar repression strength but cause lower noise on their target genes. To investigate how gene expression noise can influence the behavior of synthetic gene circuits, we further applied the results to a transcription activator-like effector repressor (TALER) switch circuit that can be employed to classify cell types using endogenous miRNAs as sensors. We found that the increase in noise mediated by miRNAs could significantly improve the classification accuracy by enhancing cell state transition. In summary, this work quantitatively characterized patterns of the miRNA-mediated modulation of gene expression noise as well as how such modulation influences the performance of synthetic gene circuits. The results provide novel insight into the function of miRNAs in gene regulatory networks and could be useful to guide the design of synthetic gene circuits to confer robustness or variability.

## Results

### Measurement of gene expression noise by a dual-fluorescent reporter system

To quantify the modulation of gene expression noise by endogenous miRNAs, we constructed a dual-fluorescent reporter system to measure the expression levels and noise of fluorescent proteins in HeLa cells. The system is composed of two fluorescent proteins (mKate2 and EYFP) that are transcribed from a bidirectional promoter. MREs are fused to the 3’ untranslated region (3’UTR) of EYFP, whereas the expression of mKate2 is not regulated by miRNAs (Fig. 1A). Therefore, the fluorescence intensity of mKate2 can be used to indicate the transcription rate of the promoter. After the transient transfection of the system into HeLa cells, the intensity of mKate2 and EYFP was quantified by flow cytometry (Fig. 1B). Cells with similar transcription rates were binned according to their mKate2 fluorescence intensity (Fig. 1B). The mean value and noise (coefficient of variation, CV) of the EYFP fluorescence intensity in each bin were calculated (Fig. 1C-E).

**Fig. 1.**
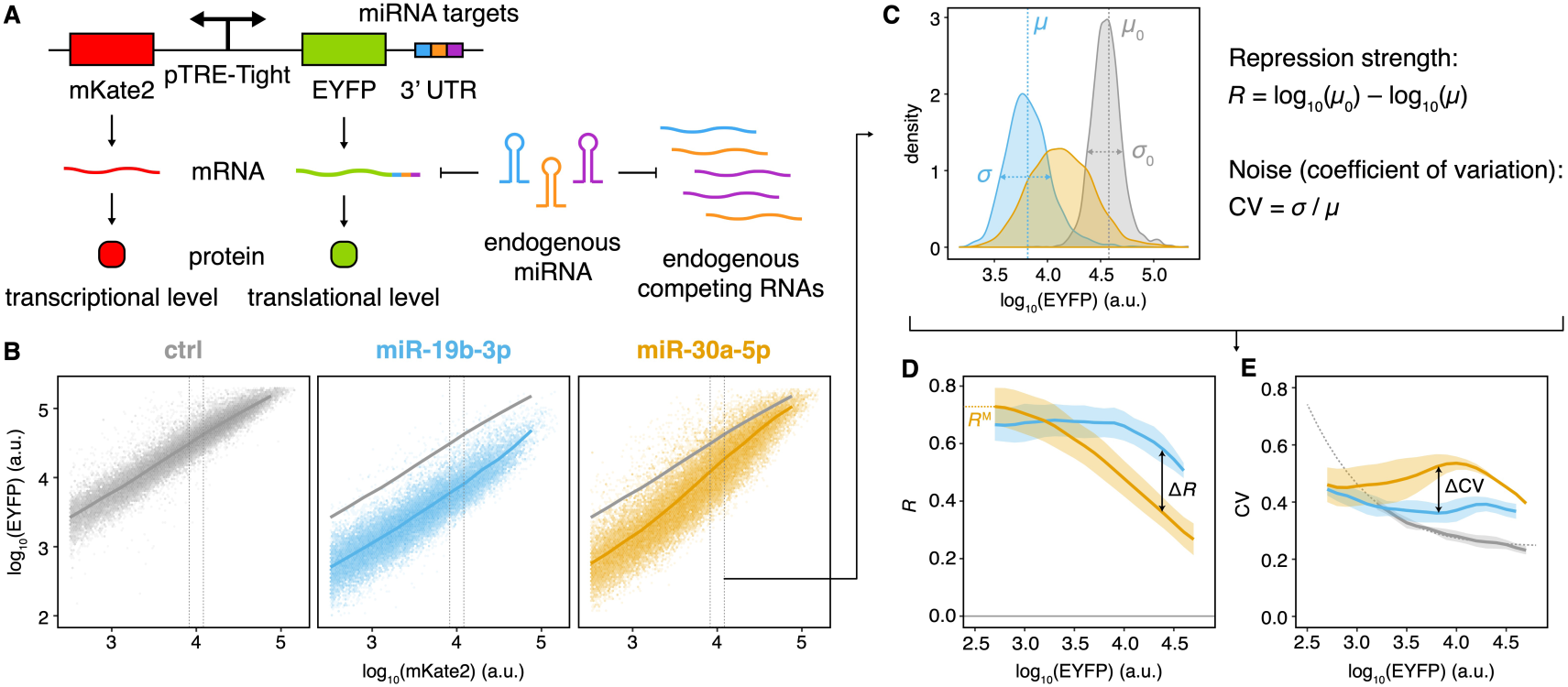
Measurement of gene expression noise by a dual-fluorescent reporter system. (*A*) Schematic diagrams of the dual-fluorescent reporter system. (*B*-*E*) Procedures for calculating gene expression noise using flow cytometry data. Flow cytometry results of systems with no MRE (gray), the single miR-19b-3p MRE (blue), or the single miR-30a-5p MRE (orange) are shown in (*B*). Cells were binned according to mKate2 fluorescent intensity (*C*). The repression strength of miRNAs (*D*) and the gene expression noise (*E*) in each bin were further calculated. The lines represent the mean value of EYFP in different mKate2 bins in (*B*). The lines and shading show the mean ± SD of three independent replicates in (*D*) and (*E*). The dotted line in (*E*) represents the fitted noise of reporters without MREs.

We identified the 20 endogenous miRNAs showing the highest expression in HeLa cells by high-throughput sequencing and cloned one perfectly complementary MRE to the 3’UTR of EYFP for each of them (Fig. S1 and Table S1). Then, we measured the correlation between the strength of miRNA-mediated repression (*R*, logarithmic fold change) and the noise of EYFP (CV) at different EYFP levels (Fig. 2A and Fig. S2). Previous studies (20) have demonstrated that the repression strength of miRNAs is a critical factor in miRNA-mediated noise control. Higher repression strength could reduce intrinsic noise, which is mainly contributed by the stochastic gene transcription, to a greater extent, thus leading to significant noise reduction at low gene expression levels. Additionally, the noise of miRNAs can propagate to the noise of the observed gene as extrinsic noise, which is dominant at high expression levels. Consistent with these findings, we found that miRNA-mediated repression strength was negatively correlated with the noise of EYFP when expressed at a low level but positively correlated under high expression (Fig. 2A). However, the correlation was not sufficiently high when gene expression levels were intermediate. Genes that are repressed to a similar extent may show quite different noise levels (Fig. 2A), leading to the speculation that other properties of miRNAs can also influence gene expression noise.

**Fig. 2.**
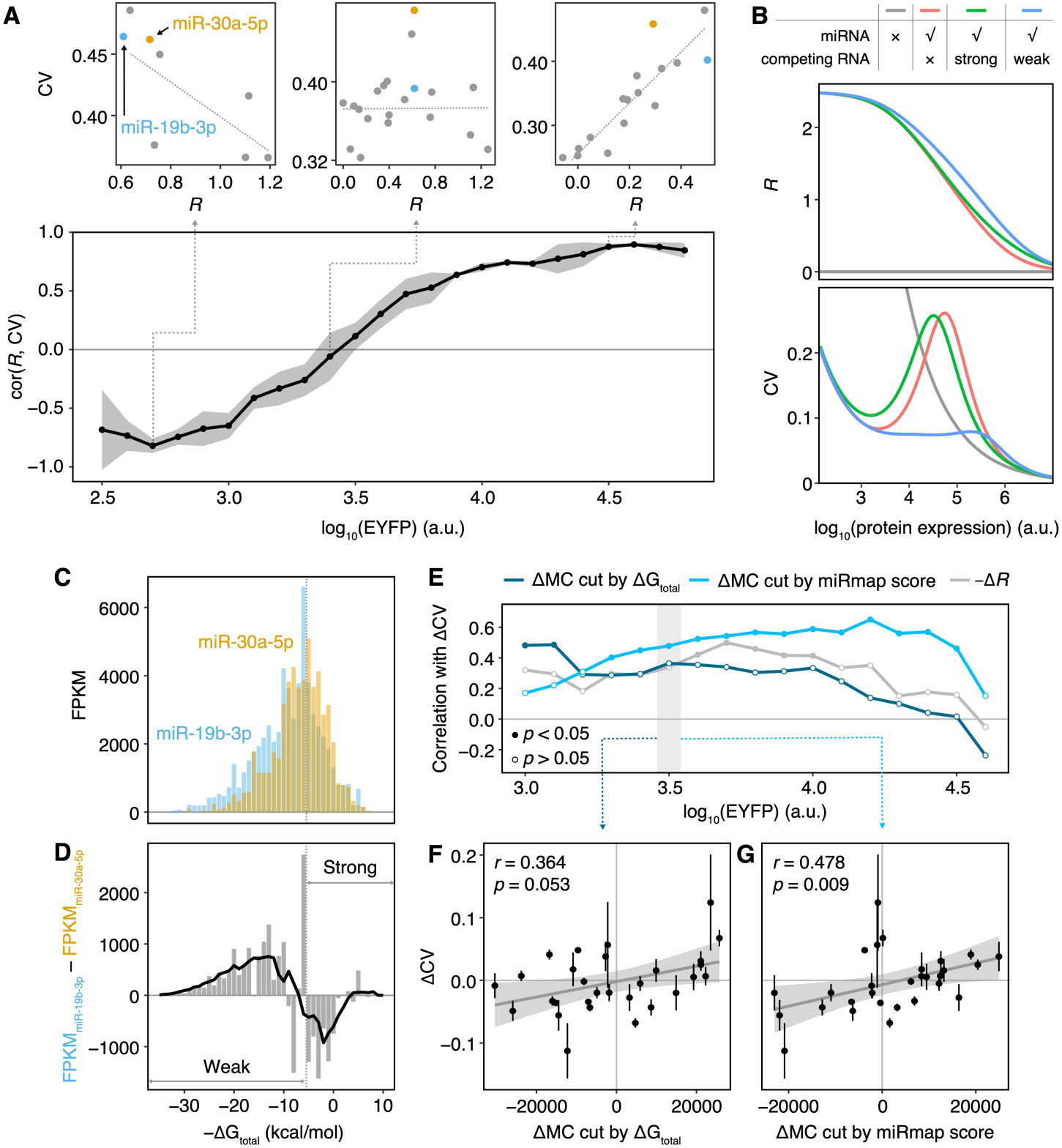
The influence of repression strength and competing RNAs on gene expression noise. (*A*) Spearman correlation between repression strength (*R*) and noise (CV) among reporter systems containing MREs with a single target of the top 20 most highly expressed miRNAs in HeLa cells at different expression levels of EYFP. Solid lines and shading represent the mean ± SD with three independent replicates. Dashed lines represent the linear regression results of points. (*B*) Simulation results of the influence of competing RNAs on gene expression levels and noise. The simulation parameters are shown in Table S3. (*C*) Competing RNA abundance associated with miR-19b-3p and miR-30a-5p under different ΔGtotal values. (*D*) The difference between competing RNA abundance associated with miR-19b-3p and miR-30a-5p under different ΔGtotal values. The black line represents the smoothing of the abundance distribution. The gray line represents the threshold of ΔGtotal that divides the competing RNAs into strong and weak groups. (*E*) Spearman correlations between ΔCV and ΔMC divided by ΔGtotal (dark blue line), between ΔCV and ΔMC divided by the miRmap score (light blue line), and between ΔCV and –Δ*R* (gray line). Solid and hollow points represent the significance of the correlation coefficients. *F-G*) Spearman correlations between ΔCV and ΔMC divided by the ΔGtotal threshold (*F*) and between ΔCV and ΔMC divided by the miRmap score threshold (*G*) at the EYFP expression level of approximately 10^3.5^ a.u. Each point represents the mean ± SD with three independent replicates. Gray lines and shading represent the linear regression and the 0.95 confidence interval of the results.

### Competing RNAs of miRNAs modulate gene expression noise

It has been shown that each miRNA targets tens or hundreds of different endogenous RNAs (28), yet most of them are only modestly repressed (less than twofold) (29–31). These RNAs compete for miRNA binding with each other, mutually playing the role of competing RNAs and constituting a complex miRNA-mediated regulatory network. By combining mathematical modeling and single-cell RNA-seq analysis, we previously showed that the abundant weak targets of miRNAs have the capacity to buffer gene expression noise (26, 27). In addition, previous research using a similar dual-fluorescent reporter system investigated how the strength of exogenous competing RNAs introduced by transfected plasmids influences gene expression noise (25). However, the binding affinities of these exogenous competing RNAs to miRNAs are much higher than those of endogenous interactions, so the results can hardly represent how endogenous miRNA-mediated regulatory networks modulate gene expression noise.

To investigate how competing RNAs regulate gene expression noise, we performed computational simulations using a minimal miRNA-competing RNA model that we and others have used previously (26, 32, 33). The model describes the interactions between one miRNA and two RNAs that can bind the miRNA competitively. The simulation results showed that when repression strengths were similar, the noise of lowly expressed genes was not sensitive to competing RNAs. However, at intermediate or high expression levels, genes targeted by miRNAs with weak competing RNA pools tended to show lower noise than genes that were not regulated by any miRNAs and those regulated by miRNAs with strong competing RNA pools (Fig. 2B).

To verify the computational results, we used a reporter system to analyze the noise modulation patterns of miRNAs with similar repression ability to exclude the influence of repression strength on noise. As the repression strength may be reduced with the increase of gene expression level (34), we took the repression strength at the lowest EYFP expression level as the maximum repression strength (*R*_M_) to describe the repression ability of a miRNA. To figure out the impact of noise on miRNAs with same repression strength, we chose all miRNA pairs exhibiting a difference of maximum repression strength (Δ*R*^M^, *R*^M^_miR-A_ – *R*^M^_miR-B_) less than 0.1 for further analysis. For example, reporters with a single perfectly complementary MRE of miR-30a-5p or miR-19b-3p showed similar repression strengths and noise levels at low expression (Fig. 1D). However, at high expression, the reporter with the miR-19b-3p MRE showed less noise than that with the miR-30a-5p MRE (Fig. 1E). We further quantified the levels of the competing RNAs of these miRNAs by RNA-seq and used the total system energy (ΔG_total_) calculated by miRmap (35) to represent the interaction strength between miRNAs and their competing RNAs. In comparison with miR-19b-3p, miR-30a-5p tended to be associated with more substantial levels of strong competing RNAs and lower levels of weak competing RNAs (Fig. 2C-D). Therefore, consistent with the computational model predictions (Fig. 2B), the different noise levels observed for this reporter pair could be partially explained by distinct interaction strength distributions of the competing RNA pool.

We performed a similar analysis for all the other pairs of reporters targeted by miRNAs with |Δ*R*^M^| < 0.1. The competing RNAs were divided into a strong group and a weak group with a ΔG_total_ threshold (7 kcal/mol), and the difference between the levels of competing RNAs in the strong group and the weak group was defined as the noise modulation capacity (MC, FPKM_strong_ – FPKM_weak_) to roughly assess the integrated influence of competing RNAs on gene expression noise. Interestingly, the differences of noise between reporter pairs (ΔCV, CV_miR-A_ – CV_miR-B_) with similar maximum repression strengths showed a moderate positive correlation with the differences of the miRNA noise modulation capacity (ΔMC, MC_miR-A_ – MC_miR-B_) (Fig. 2E-F). This positive correlation was even more significant when using the miRmap score to define the interaction strength (miRmap score threshold = –0.05) (Fig. 2E and G). These observations supported the hypothesis that gene expression noise is buffered by weak competing RNAs.

### The composition of MREs influences miRNA-mediated noise modulation

The 3’UTRs of both endogenous genes and genes included in synthetic circuits often contain many miRNA binding sites (36, 37). Some 3’UTRs are targeted by the same miRNA multiple times to enhance the repression strength of miRNAs (18). Some 3’UTRs are targeted by different miRNAs, constituting a complex regulatory network controlling various biological processes robustly and precisely (38). Therefore, it is necessary to investigate how the composition of MREs can influence gene expression noise.

To investigate how multiple MREs of the same miRNA can modulate noise, we constructed reporters that contain three perfect complementary tandem MREs for each of the 20 miRNAs showing the highest expression in HeLa cells (Fig. 3A and Fig. S3). Compared to reporters with a single MRE, reporters with triple MREs showed a greater repression strength at low expression levels, but the repression strength decreased markedly as expression increased (Fig. 3C), which is a consequence of the saturation effect of miRNA regulation (34). The sensitivity of the strength of gene expression repression (d*R*) around the threshold generated by saturation therefore increases in reporters with triple MREs (Fig. 3E), which has been shown to increase gene expression noise in previous studies (14, 26). Therefore, reporters with triple MREs showed stronger noise reduction at low expression levels in comparison with those with a single MRE but exhibited a remarkable increase in noise at high expression levels (Fig. 3I). Furthermore, accompanied by the alteration of the threshold, the regime with maximum sensitivity showed a shift to lower expression levels in triple-MRE reporters, causing an increase in noise even at low expression levels, which counteracted the noise reduction arising from the increased repression strength of triple MREs (Fig. 3G and I). This phenomenon is consistent with previous work (25, 34).

**Fig. 3.**
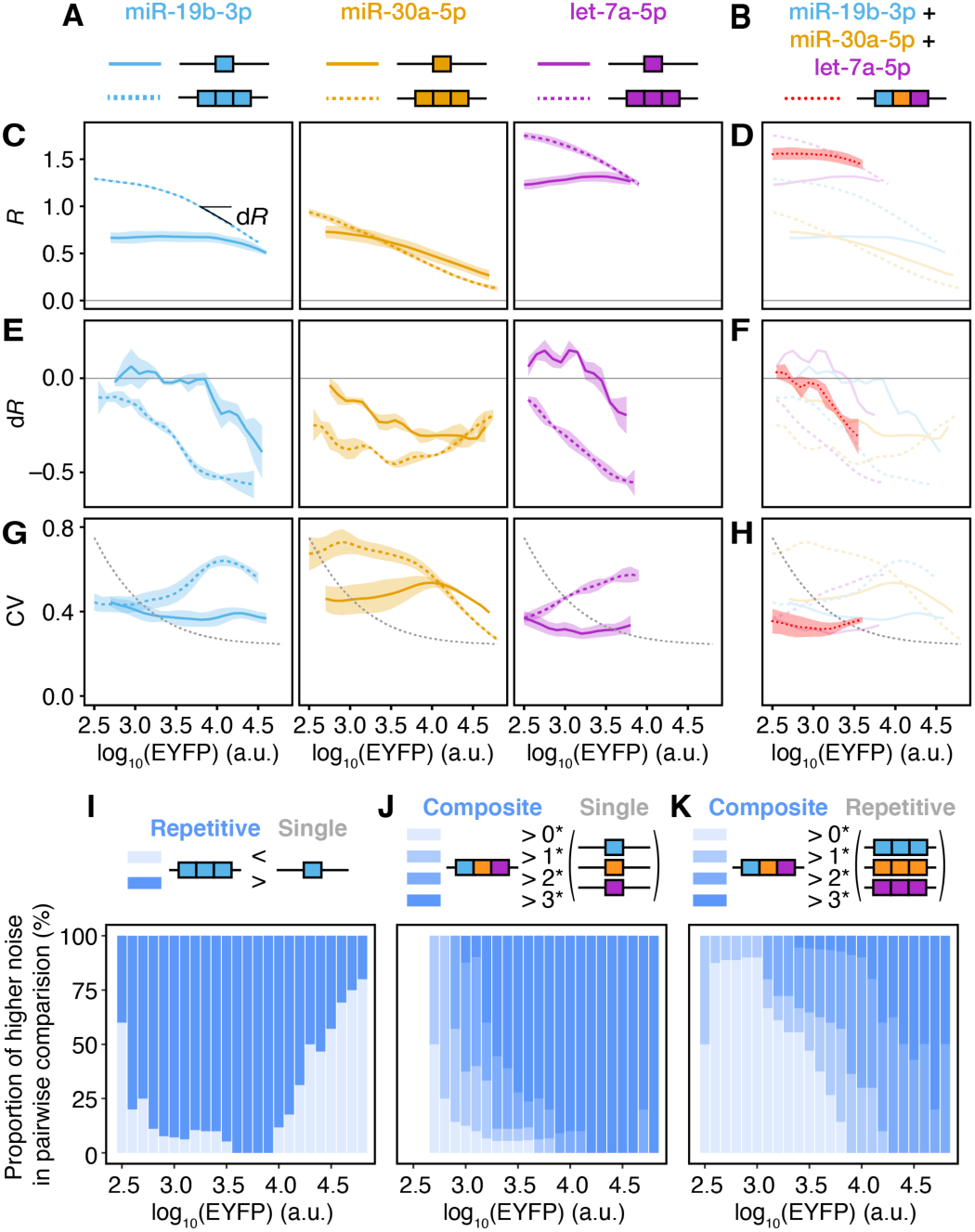
The influence of MRE composition on gene expression noise. (*A-B*) Schematic diagrams of a single MRE (*A*), triple repetitive MREs (*A*), and composite MREs (*B*). (*C*-*H*) Repression strength (*C-D*), saturation (*E-F*, the difference of repression strength) and noise (*G-H*) of reporters with a single MRE (colored solid lines), triple MREs (colored dashed lines), and composite MREs (red dotted lines). Lines and shading represent the mean ± SD with three independent replicates. Gray lines represent reporters without the regulation of miRNAs. (*I-K*) Comparison of noise (*I*) between the reporter with triple MREs and the reporter with the corresponding single MRE, (*J*) between the reporter with composite MREs and all three reporters with the corresponding single MRE, and (*K*) between the reporter with composite MREs and all three reporters with the corresponding triple MREs. Only EYFP expression levels measured in ≥ five comparison groups are shown.

We further used the reporter systems to assess the modulation pattern of MREs composed of targets of different miRNAs. Each composite MRE is composed of three perfectly complementary MREs that are targeted by miRNAs with adjacent expression level ranks in HeLa cells. The top 20 most highly expressed miRNAs in HeLa cells were selected, and they constituted 18 sequential composite targets. Interestingly, the composite MREs exhibited distinct noise modulation patterns compared to repetitive MREs and a single MRE (Fig. 3B, S4, and S5). In comparison with reporters with a single MRE, reporters with composite MREs showed higher miRNA-mediated repression strength and thus lower noise at low expression levels (Fig. 3D, H, and J). Additionally, the composite MREs could lead to greater saturation (Fig. 3F), thus increasing noise at very high expression levels (Fig. 3H, J, and S4). Compared with triple MREs, the composite MREs showed similar repression levels but a higher saturation threshold, leading to a wide range of noise reduction levels compared with repetitive MREs except in the case of extremely high expression levels (Fig. 3H and K and S5).

### MiRNA-mediated noise modulation could enhance state transition and improve the accuracy of synthetic cell-type classifiers

miRNAs have been widely employed in synthetic gene circuits as input signals to classify cells of different types or in different states (39–44). Previous studies have shown that gene expression noise can impact the performance of synthetic gene circuits (3, 14), so it is reasonable to hypothesize that the ability of miRNAs to control gene expression noise could further modulate the behaviors of gene circuits with miRNAs as inputs. Here, we used a well-established miRNA-mediated TALER circuit that behaves as a cell-type classifier (42) (Fig. 4A and S6A) to investigate whether miRNA-mediated noise modulation can affect the classification accuracy of the circuit.

**Fig. 4.**
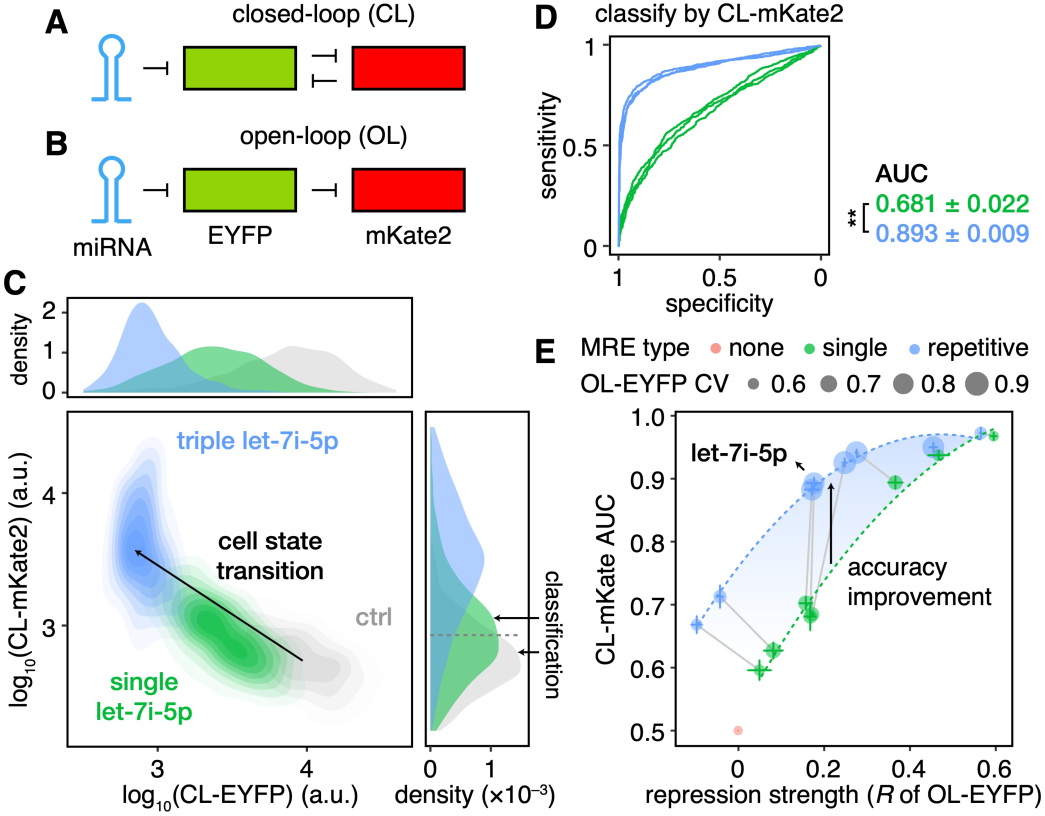
miRNAs can regulate the behavior of TALER switches by modulating gene expression noise. (*A*) Schematic diagrams of closed-loop (CL) TALER switches. (*B*) Schematic diagrams of open-loop (OL) TALER circuits. (*C*) Joint and marginal distribution of EYFP and mKate2 in CL switches. (*D*) ROC curves obtained when classifying cells by mKate2 in CL switches. Student’s *t*-tests were performed between the AUCs of two groups, and the significance levels are indicated by **: *p* < 0.01. (*E*) The relationship between the repression strength of MREs in OL circuits and the AUCs classified by mKate2 in CL switches. The size of the point represents the noise level (CV) of EYFP mediated by the MRE in OL circuits. Dashed curves represent the loess regression of the points. Each point represents the mean ± SD with three independent replicates. Points that represent reporters with single MRE or triple MREs of the same miRNA are connected by gray lines.

As shown in Figure 4A and S6A, the TALER switch was composed of two TALER genes that could repress the transcription of each other, forming a closed-loop (CL) topology. The TALER genes were fused with EYFP or mKate2 so that their expression levels were reflected by the intensity of these fluorescent proteins. The expression of both TALERs was driven by the transcription activator Gal4VP16 fused with TagBFP. We selected the top eight most highly expressed miRNAs in HeLa cells and cloned one perfectly complementary MRE or triple tandem perfectly complementary MREs for each of them to the 3’UTR of EYFP-TALER. To exclude the influence of the transfection efficiency on EYFP and mKate2 intensity, we binned cells according to their TagBFP intensity and chose cells with TagBFP intensity between 10^4.3^ and 10^4.4^a.u. for further analysis. The results for other bins exhibited similar trends and are shown in supplementary figure S7.

Previous studies (42) have shown that CL switches without miRNA regulation exhibit a state with moderate levels of both EYFP and mKate2. When the expression of EYFP is repressed by miRNA, CL switches will transition to a low-EYFP, high-mKate2 state. Cell state transitions were successfully observed in this study, but the extent of the transition was quite different across these CL switches with different MREs (Fig. 4C). Some CL switches exhibited an approximately complete transition to the low-EYFP, high-mKate2 state, while others exhibited intermediate states. The CL TALER switches could be used to classify different cell types or states employing different miRNAs as input (42, 45), and an incomplete transition will decrease the classification accuracy between the circuits with and without the inputs (Fig. 4C). We depicted the classification accuracy in a receiver operating characteristic (ROC) curve and quantified the accuracy by using the area under the ROC curve (AUC) (Fig. 4D). Furthermore, to determine the repression-strength-independent influence of noise modulation mediated by miRNAs on the performance of CL switches, we also quantified the repression strength of each MRE by using an open-loop (OL) TALER circuit, the only difference of which in comparison with the CL switch was that mKate2-TALER could not repress the expression of EYFP-TALER (Fig. 4B and S6B).

As shown in Figure 4E, the classification accuracies were positively correlated with the repression strength. However, circuits with MREs sharing similar repression strengths in OL circuits could exhibit distinct AUCs in CL switches. In particular, CL switches with triple tandem MREs (blue points) showed higher classification accuracies than those with a single MRE (green points), which was consistent across all TagBFP intensity bins (Fig. S7). For instance, single let-7i-5p MRE and triple let-7i-5p MREs could lead to similar repression strengths in OL circuits (Fig. 4E and S8A), but only the CL switch with triple let-7i-5p MREs exhibited a near-complete state transition (Fig. 4C) in comparison with the CL switch with single let-7i-5p MRE, leading to higher classification accuracy (Fig. 4D-E).

To understand the mechanism of such phenomena, we examined the behavior of OL circuits with these MREs. The OL circuits can be regarded as a signal transduction cascade in which the signal of miRNA propagates to mKate2-TALER via EYFP-TALER. We found that although similar repression strengths in the first step (EYFP) were exerted, circuits with triple MREs could induce a greater expression increase in the second step (mKate2) (Fig. S8B). For example, although the OL circuit with triple let-7i-5p MREs exhibited the same EYFP level as that with a single let-7i-5p MRE, the former circuit showed a significantly higher mKate2 level than the latter (Fig. S8A). The experimental results of the dual-fluorescent reporter system showed that triple MREs could introduce higher noise at low and intermediate expression levels in comparison with a single MRE, independent of repression strength (Fig. 3). Therefore, we speculated that the miRNA-mediated modulation of noise may affect the signal transduction efficiency of the OL circuits, and it may further influence the classification accuracy of CL switches.

We performed Monte Carlo simulations (46) to investigate the impact of noise. In OL circuits, the results showed that when mean EYFP levels were equal, higher EYFP noise could lead to a higher mKate2 level, resulting from the nonlinear relationship between EYFP and mKate2 (Fig. S9A-B). In CL switches, the increase in the mKate2 level caused by higher EYFP noise could in turn suppress the expression of EYFP, thus bringing about a significant change of the distribution to a lower-EYFP and higher-mKate2 situation (Fig. S9D-F). Besides, in CL switches, where the expression of EYFP was repressed by mKate2, the stronger saturation effects of repetitive MREs in comparison with the single MRE (Fig. 3E) could lead to a more remarkable increase of repression strength in CL switches (Fig. S9C), which resulted in the alteration of the steady state to a low-EYFP and high-mKate2 state (Fig. S9D-E). Therefore, the alteration of repression strength caused by saturation promoted the cell state transitions in CL switches, and the high expression noise modulated by MREs further enhanced the cell state transitions (Fig. S9E and F), both of which upheld the improvement of classification accuracy (Fig. S9G). The results suggested that detailed characteristics of miRNA regulation should be taken into account in the design of synthetic circuits with miRNAs.

## Discussion

We applied mathematical modeling and a dual-fluorescent reporter system to investigate how competing RNAs and miRNA binding sites of genes influence the expression noise of genes at different expression levels. We showed that genes targeted by miRNAs with weak competing RNAs tend to exhibit lower noise than those without MREs or those targeted by miRNAs with strong competing RNAs at a wide range of gene expression levels. In addition, repetitive MREs of the same miRNA can reduce noise at low expression levels but increase gene expression noise at high expression levels compared to a single MRE, while composite MREs composed of targets for different miRNAs can maintain low-expression noise reduction but reduce the increase of high-expression noise. We showed that the proper choice of miRNAs and corresponding MREs to modulate gene expression noise could significantly enhance cell state transition and improve the performance of synthetic gene circuits for cell classification.

Each miRNA has tens or hundreds of targets, but most of them are weakly repressed, raising questions about the function of the widespread weak interactions between miRNAs and their targets (29–31). A generally accepted hypothesis posited that miRNAs usually make fine-scale adjustments to most of their targets (17, 19, 28). An alternative proposed view is that a large proportion of miRNA targets are competitive inhibitors of miRNAs. Moderate repression of these targets might not lead to consequences at the physiological level (28) but may function to titrate miRNA activity, instead of being regulated by miRNAs (17). Our results suggested that widespread weak interactions might act as buffers to reduce gene expression noise (Fig. 2) and thus confer robustness to gene regulatory networks, similar to previous theoretical analysis (47, 48) and our observations at the mRNA level (27).

It is noteworthy that current algorithms for predicting miRNA targets, such as miRmap (35, 49), TargetScan (50), and PITA (51), were designed to discover targets that are thought to be repressed by corresponding miRNAs with high confidence. To reduce the false-positive rate, these algorithms usually ignore weak or nonconserved targets. However, it has been suggested that the number of these weak targets that cannot be predicted by current algorithms is quite large and that they might play important roles in the regulation of gene expression by miRNAs (52). According to our computational simulation results, these widespread weak targets can buffer gene expression noise. Therefore, if algorithms for predicting these weak targets can be developed in the future, we could better understand the modulation of gene expression noise by miRNAs and employ the results of this study to explain the function of endogenous miRNAs in biological processes or modulate the performance of synthetic gene circuits.

In synthetic gene circuits with miRNAs as inputs, MREs are often designed as tandem repeats of the same miRNA instead of a single MRE of the miRNA (39, 42, 53). We have shown that genes with repetitive MREs may exhibit higher noise levels at high expression levels or even globally than those with a single MRE, demonstrating that the composition of MREs can strongly influence gene expression noise. In addition, as shown in the TALER-based cell-type classifiers, switches with repetitive MREs exhibited better classification accuracies than those with a single MRE when subjected to similar repression strengths measured in OL circuits. The results indicated that during the design of MREs in synthetic gene circuits, it is necessary to consider not only the repression strengths measured in some certain conditions, but also miRNA-mediated modulations of noise and saturation as well as the intricate behaviors of the modulations in gene regulatory networks.

The engineering of large-scale synthetic gene circuits is limited by the poor stability of the complex system, which is usually caused by noise in gene expression. Previous studies have revealed several strategies for modulating gene expression noise without changing the mean expression levels, including the mutation of TATA boxes (2, 54), the regulation of transcriptional rates and translational rates at the same time (3), the alteration of epigenetics (55) and the arrangement of two transcriptional regulators (56). Our investigation of the influence of miRNA properties on gene expression noise might provide a new tool for modulating noise in synthetic gene circuits and thus regulating their behaviors.

In summary, we performed an elaborate analysis of how competing RNAs and the composition of MREs influence gene expression noise and how such modulation could further impact the performance of synthetic gene circuits. The results provided an explanation for the function of the widespread weak interactions between miRNAs and their targets and presented guidelines for the design of miRNA binding sites in synthetic gene circuits to better employ miRNAs as endogenous signals.

## Materials and Methods

### Reagents and enzymes

Polynucleotide kinase (PNK), ATP, T4 ligase, and all restriction endonucleases were purchased from New England Biolabs. Oligonucleotides were synthesized by Genewiz. Doxycycline (Dox) was purchased from Clontech.

### Construction of dual-fluorescent reporter systems and TALER switches

The sequences of the single MREs are listed in Table S1. The sequences of the triple MREs consisted of three tandem single MRE sequences without gaps. The sequences of the composite MREs were joint by single MRE sequences of three miRNAs with adjacent ranks of expression levels without gaps. The MREs were synthesized as oligonucleotides, which were annealed and then phosphorylated using PNK and ATP for ligation. The MREs were then inserted into the 3’UTR of EYFP in the dual-fluorescent reporter system via the SpeI and HindIII digestion sites with T4 ligase.

The TALER-based switches were modified from circuits described in a previous study (42). The closed-loop (CL) switch (Fig. S6A) was composed of three plasmids. The constitutively expressed transcription activator Gal4VP16, which was fused with TagBFP via a self-cleaving 2A linker (Gal4VP16-2A-TagBFP, or Gal4VP16-TagBFP in short) (57), drove the expression of EYFP-2A-TALER9 and mKate2-2A-TALER14. The promoter of EYFP-TALER9 was flanked with four TALER14 binding sites (T14), and the promoter of mKate2-TALER14 was flanked with four TALER9 binding sites (T9). Therefore, EYFP-TALER9 and mKate2-TALER14 exerted mutual expression inhibition against each other. The MREs were inserted into the 3’UTR of EYFP-TALER9 via the SpeI and MluI digestion sites with T4 ligase. Compared with the CL switch, the only difference in the open-loop (OL) circuit was that mKate2-TALER14 was replaced by mKate2-TALER10 so that the expression of mKate2-TALER10 could not inhibit the expression of EYFP-TALER9 (Fig. S6B).

### Cell culture, transient transfection and flow cytometry

HeLa cells (originally obtained from ATCC) were grown in DMEM with 4.5 g/L glucose (Gibco) supplemented with 10% FBS (Gibco) at 37 °C and 5% CO_2_. On the day before transfection, approximately 1.6×10^5^ HeLa cells were plated in 12-well plates. Lipofectamine LTX (Thermo Fisher) was used for transfection according to the manufacturer’s protocol. For the observation of dual-fluorescent reporter systems, transfection with 40 ng of the plasmids carrying the dual-fluorescent reporter, 40 ng of the plasmids carrying the reverse tetracycline-controlled transactivator (rtTA) gene, and 420 ng of pDT004, a plasmid with no protein-coding sequences (42), was performed in the wells of a 12-well plate. For the observation of TALER switches, the transfection of 100 ng of the plasmids carrying Gal4VP16-TagBFP, 100 ng of the plasmids carrying EYFP-TALER9-MRE, and 100 ng of the plasmids carrying mKate2-TALER10 (for OL) or mKate2-TALER14 (for CL) in the wells of a 12-well plate. On the day of transfection and one day after transfection, the medium was refreshed with DMEM. For the dual-fluorescent reporters, we supplemented DMEM with 1 μg/ml Dox to induce their expression. Cells were harvested 48 h after transfection and centrifuged at 500 g for 3 min at room temperature. Then, the cells were washed with PBS once and resuspended in PBS. Flow cytometry was conducted using Fortessa flow analyzers (BD Biosciences) with the settings listed in Table S2. For the dual-fluorescent reporter systems, approximately 3×10^4^ mKate2-positive cells were collected. For the TALER switches, approximately 5×10^4^ TagBFP-positive cells were collected.

### High-throughput sequencing of HeLa cells

The miRNA-seq of HeLa cells was conducted on the HiSeq 2500 platform (Illumina) and analyzed by using miRDeep2 (v 0.1.2). The RNA-seq of HeLa cells was conducted on the NovaSeq 6000 platform (Illumina) and analyzed by using TopHat (v 2.1.1) and HTSeq (v 0.6.1).

### Flow cytometry data processing

All data processing and analysis described below were performed by using R v3.6.1. For the dual-fluorescent reporter system used to measure gene expression noise, the data were processed as previously described with some modifications (20), as follows. Cells were binned according to the mKate2 intensity (bin width 0.2 in log10 space). Cells with mKate2 intensity of less than 10^2^ were considered to be untransfected cells and were thus removed. In each bin, cells above the 0.05 quantile and below the 0.95 quantile of the EYFP intensity distribution were selected. Three independent biological replicates were performed for each reporter. The noise of reporters without MREs was fitted using the method described in Ref. (20).

For the data of TALER circuits, the cells were binned according to the TagBFP intensity (bin width 0.1 in log10 space) to control the transfection efficiency. Cell type classification was performed between the CL switches with MREs and without MREs in the same replicate.

The repression strength, *R*, of each miRNA was calculated as the difference between the logarithmic mean value of EYFP without miRNA regulation (*μ*_0_) and with miRNA regulation (*μ*) at the same transcriptional level (in the same mKate2 bin for the dual-fluorescent reporter system or the same TagBFP bin for the OL TALER circuits): *R* = log_10_(*μ*_0_) – log_10_(*μ*).

For the dual-fluorescent reporter system, *R* and CV were interpolated in the space of the EYFP mean expression level. The repression strength at the lowest EYFP expression level was regarded as the maximum repression strength, *R*^M^. miRNA pairs with similar *R*^M^ (|*R*^M^_miR-A_ – *R*^M^_miR-B_| < 0.1) values were considered to be pairs of miRNAs with the same repression strength for further analysis in Fig. 2E-G.

### Characterization of competing RNAs

We calculated competing RNA abundance from the RNA-seq data of HeLa cells and the miRNA target predictions for humans by using miRmap (release mirmap201301e). For the analysis based on ΔG_total_, all transcripts with ΔG_total_ < 7 kcal/mol were regarded as strong competing RNAs; otherwise, they were regarded as weak competing RNAs. The total abundance (FPKM) of the strong and weak competing RNAs was used to define the noise modulation capacity (MC): MC = FPKM_strong_ – FPKM_weak_. For analysis based on the miRmap score, all transcripts with a miRmap score < –0.05 were regarded as strong competing RNAs.

### Mathematical modeling of gene expression noise

The mathematical model and parameter settings for simulating the noise modulation of competing RNAs were adopted from our previous work (26). The model is briefly described using ordinary differential equations (ODEs) as follows:

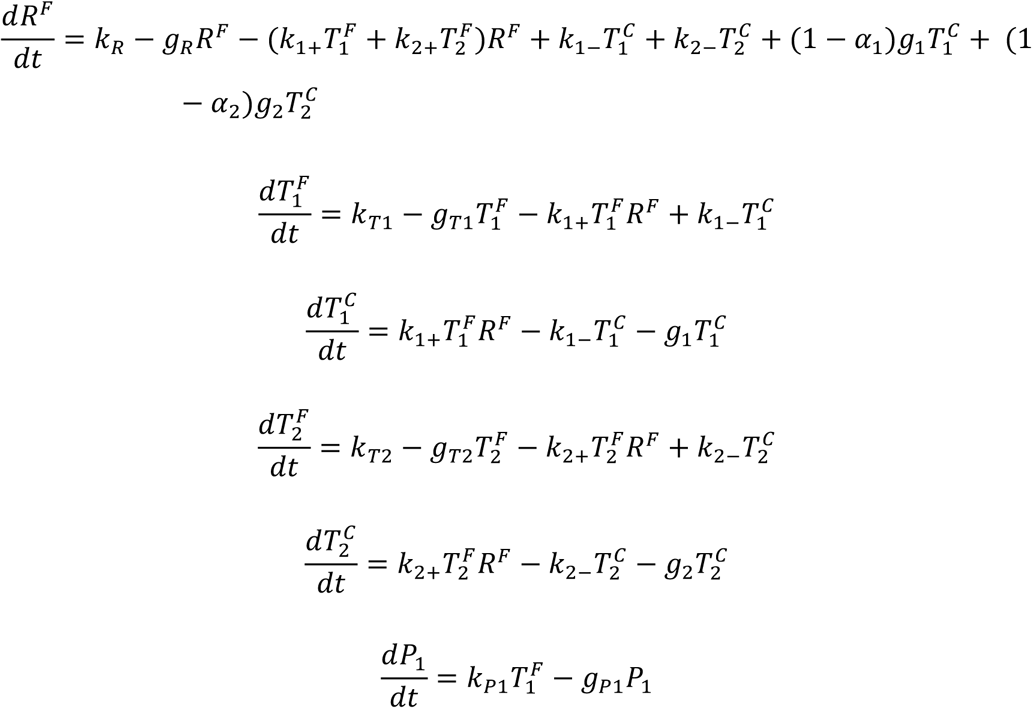

Where *R*^F^ denotes the concentration of free miRNA; 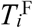 denotes the concentration of free RNA #*i*; 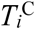 denotes the concentration of the complex of miRNA and RNA #*i*; *P*_1_ denotes the protein of RNA #1. In this model, RNA #1 represents the RNA transcribed from the observed gene, and RNA #2 represents the competing RNA. *k*_*Ti*_ is the transcription rate of RNA #*i. g*_*Ti*_ is the degradation rate of RNA #*i*. RNA #*i* and the miRNA form a complex at a rate of *k*_*i*+_, and the complex dissociates at a rate of *k*_*i*–_ and degrade at a rate of *g*_*i*_. When complex #*i* degrades, miRNAs on the complex degrades with a probability of *α*_*i*_. The production and degradation rate of the protein of RNA #1 are *k*_*P*1_ and *g*_*P*1_ respectively. The transcription and degradation rate of the miRNA are *k*_*R*_ and *g*_*R*_ respectively. The noise levels of all the molecular species are calculated using fluctuation-dissipation theorem as in Ref. (26). The parameters for Fig. 2B are shown in Table S3.

### Stochastic simulation of TALER circuits

To model how the noise of EYFP-TALER influences the level of mKate2-TALER, we conducted a stochastic simulation using the Gillespie algorithm (46). The EYFP-TALER was denoted as *E*, and the mKate2-TALER was denoted as *K*. The transcription of *K* mRNAs (*T*_*K*_) was repressed by *E* proteins (*P*_*E*_) in both OL and CL circuits, while the transcription of *E* mRNAs (*T*_*E*_) was repressed by *K* proteins (*P*_*K*_) only in CL switches. *T*_*E*_ and *T*_*K*_ transcribes at a rate of *k*_*TE*_ and *k*_*TK*_ respectively without repression. The repressions of *P*_*K*_ for *T*_*E*_ and *P*_*E*_ for *T*_*K*_ follow the Hill’s function. For the repression of *P*_*E*_ for *T*_*K*_, *K*_*E*_ denotes the number of *P*_*E*_ that gives 50% repression of *T*_*K*_, and *m* denotes the Hill coefficient of this reaction. For the repression of *P*_*K*_ for *T*_*E*_, which only exists in CL switches, *K*_*K*_ denotes the number of *P*_*K*_ that gives 50% repression of *T*_*E*_, and *n* denotes the Hill coefficient of this reaction. The values of Hill coefficients were adopted from Ref (42). *g*_*TE*_, *g*_*TK*_, *g*_*PE*_ and *g*_*PK*_ denote the degradation rate of *T*_*E*_, *T*_*K*_, *P*_*E*_ and *P*_*K*_ respectively. The deterministic model of OL circuits is described in the following ODEs:

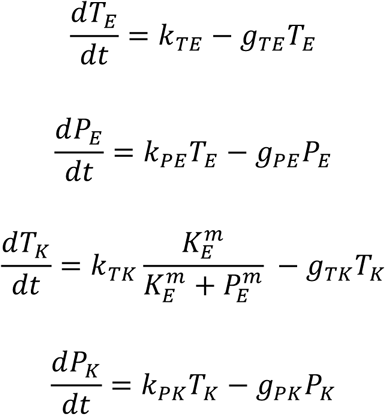

In CL switches, the first equation is replaced by

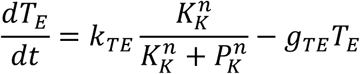

Two major factors of miRNA regulation, repression strength and gene expression noise, were simulated separately. We defined *k*_*TE*0_ and *g*_*TE*0_ as the production rate and degradation rate of *T*_*E*_ in the condition without the regulation of miRNA. Furthermore, we defined the relative repression rate *R* and the turnover rate *t* for a certain condition, where

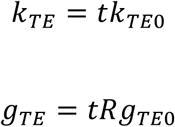

The relative repression rate *R* represents the condition with or without the regulation of miRNA and the difference of repression strengths in CL switches. The turnover rate *t* represents different expression noise of EYFP-TALER. A high turnover rate indicates low noise, and a low turnover rate indicates high noise. The parameters are listed in Table S4.

For simulation using Gillespie algorithm, there are 8 reactions total: the production and degradation of *T*_*E*_, *T*_*K*_, *P*_*E*_, *P*_*K*_ respectively. Their propensity functions in the Gillespie algorithm are the same with the coefficients in the ODEs. We simulated 100 trajectories for each parameter setting. After the fluctuation reached an approximately steady state (*t* = 10000 s), a total of 400 points were sampled per 100 seconds from every trajectory for plotting Fig. S9A and S9D. The mean values of EYFP and mKate2 from 10000 s to 50000 s of all trajectories were used to perform the Mann–Whitney *U* test in Fig. S9B and S9F.

## Acknowledgments

This work has been supported by the National Natural Science Foundation of China (No. 61773230, 61721003).

## Author Contributions

X.W., L.W. and S.L. designed the research; L.W. and S.L. performed the experiments and analyzed the data; L.W., S.L. and T.H. built mathematical models and performed the simulations; L.W. and S.L. wrote the manuscript, with all authors contributing to the writing and providing feedback.

## References

1. M. B. Elowitz, Stochastic Gene Expression in a Single Cell. Science 297, 1183–1186 (2002).

2. W. J. Blake, et al., Phenotypic Consequences of Promoter-Mediated Transcriptional Noise. Mol. Cell 24, 853–865 (2006).

3. H. Maamar, A. Raj, D. Dubnau, Noise in Gene Expression Determines Cell Fate in Bacillus subtilis. Science 317, 526–529 (2007).

4. G. M. Suel, R. P. Kulkarni, J. Dworkin, J. Garcia-Ojalvo, M. B. Elowitz, Tunability and Noise Dependence in Differentiation Dynamics. Science 315, 1716–1719 (2007).

5. T. Çaėatay, M. Turcotte, M. B. Elowitz, J. Garcia-Ojalvo, G. M. Süel, Architecture-Dependent Noise Discriminates Functionally Analogous Differentiation Circuits. Cell 139, 512–522 (2009).

6. H. H. Chang, M. Hemberg, M. Barahona, D. E. Ingber, S. Huang, Transcriptome-wide noise controls lineage choice in mammalian progenitor cells. Nature 453, 544–547 (2008).

7. T. Kalmar, et al., Regulated Fluctuations in Nanog Expression Mediate Cell Fate Decisions in Embryonic Stem Cells. PLoS Biol. 7, e1000149 (2009).

8. J. Hanna, et al., Direct cell reprogramming is a stochastic process amenable to acceleration. Nature 462, 595–601 (2009).

9. S. L. Spencer, S. Gaudet, J. G. Albeck, J. M. Burke, P. K. Sorger, Non-genetic origins of cell-to-cell variability in TRAIL-induced apoptosis. Nature 459, 428–432 (2009).

10. P. B. Gupta, et al., Stochastic State Transitions Give Rise to Phenotypic Equilibrium in Populations of Cancer Cells. Cell 146, 633–644 (2011).

11. A. Brock, H. Chang, S. Huang, Non-genetic heterogeneity — a mutation-independent driving force for the somatic evolution of tumours. Nat. Rev. Genet. 10, 336–342 (2009).

12. S. Hooshangi, S. Thiberge, R. Weiss, Ultrasensitivity and noise propagation in a synthetic transcriptional cascade. Proc. Natl. Acad. Sci. 102, 3581–3586 (2005).

13. J. M. Pedraza, Noise Propagation in Gene Networks. Science 307, 1965–1969 (2005).

14. W. J. Blake, M. KÆrn, C. R. Cantor, J. J. Collins, Noise in eukaryotic gene expression. Nature 422, 633–637 (2003).

15. M. Kærn, T. C. Elston, W. J. Blake, J. J. Collins, Stochasticity in gene expression: from theories to phenotypes. Nat. Rev. Genet. 6, 451–464 (2005).

16. A. Eldar, M. B. Elowitz, Functional roles for noise in genetic circuits. Nature 467, 167–173 (2010).

17. S. L. Ameres, P. D. Zamore, Diversifying microRNA sequence and function. Nat. Rev. Mol. Cell Biol. 14, 475–488 (2013).

18. D. P. Bartel, Metazoan MicroRNAs. Cell 173, 20–51 (2018).

19. D. P. Bartel, C.-Z. Chen, Micromanagers of gene expression: the potentially widespread influence of metazoan microRNAs. 5.

20. J. M. Schmiedel, et al., MicroRNA control of protein expression noise. Science 348, 128–132 (2015).

21. M. S. Ebert, P. A. Sharp, Roles for MicroRNAs in Conferring Robustness to Biological Processes. Cell 149, 515–524 (2012).

22. D. M. Kasper, et al., MicroRNAs Establish Uniform Traits during the Architecture of Vertebrate Embryos. Dev. Cell 40, 552-565.e5 (2017).

23. E. Ferro, C. Enrico Bena, S. Grigolon, C. Bosia, microRNA-mediated noise processing in cells: A fight or a game? Comput. Struct. Biotechnol. J. 18, 642–649 (2020).

24. J. Schmiedel, D. S. Marks, B. Lehner, N. Bluthgen, Noise control is a primary function of microRNAs and post-transcriptional regulation. Biorxiv (2017).

25. C. Bosia, et al., RNAs competing for microRNAs mutually influence their fluctuations in a highly non-linear microRNA-dependent manner in single cells. Genome Biol. 18 (2017).

26. L. Wei, et al., Regulation by competition: a hidden layer of gene regulatory network. Quant. Biol. 7, 110–121 (2019).

27. T. Hu, et al., Single cell transcriptomes reveal characteristics of miRNA in gene expression noise reduction. bioRxiv, 465518 (2018).

28. H. Seitz, Redefining MicroRNA Targets. Curr. Biol. 19, 870–873 (2009).

29. M. Selbach, et al., Widespread changes in protein synthesis induced by microRNAs. Nature 455, 58–63 (2008).

30. D. Baek, et al., The impact of microRNAs on protein output. Nature 455, 64–71 (2008).

31. A. Grimson, et al., MicroRNA Targeting Specificity in Mammals: Determinants beyond Seed Pairing. Mol. Cell 27, 91–105 (2007).

32. Y. Yuan, et al., Model-guided quantitative analysis of microRNA-mediated regulation on competing endogenous RNAs using a synthetic gene circuit. Proc. Natl. Acad. Sci. 112, 3158–3163 (2015).

33. J. Noorbakhsh, A. H. Lang, P. Mehta, Intrinsic Noise of microRNA-Regulated Genes and the ceRNA Hypothesis. PLoS ONE 8, e72676 (2013).

34. S. Mukherji, et al., MicroRNAs can generate thresholds in target gene expression. Nat. Genet. 43, 854–859 (2011).

35. C. E. Vejnar, E. M. Zdobnov, miRmap: Comprehensive prediction of microRNA target repression strength. Nucleic Acids Res. 40, 11673–11683 (2012).

36. R. C. Friedman, K. K.-H. Farh, C. B. Burge, D. P. Bartel, Most mammalian mRNAs are conserved targets of microRNAs. Genome Res. 19, 92–105 (2008).

37. L. S. Hon, Z. Zhang, The roles of binding site arrangement and combinatorial targeting in microRNA repression of gene expression. Genome Biol. 8, R166 (2007).

38. Z. Liufu, et al., Redundant and incoherent regulations of multiple phenotypes suggest microRNAs’ role in stability control. Genome Res. 27, 1665–1673 (2017).

39. Z. Xie, L. Wroblewska, L. Prochazka, R. Weiss, Y. Benenson, Multi-Input RNAi-Based Logic Circuit for Identification of Specific Cancer Cells. Science 333, 1307–1311 (2011).

40. K. Miki, et al., Efficient Detection and Purification of Cell Populations Using Synthetic MicroRNA Switches. Cell Stem Cell 16, 699–711 (2015).

41. B. D. Brown, et al., Endogenous microRNA can be broadly exploited to regulate transgene expression according to tissue, lineage and differentiation state. Nat. Biotechnol. 25, 1457–1467 (2007).

42. Y. Li, et al., Modular construction of mammalian gene circuits using TALE transcriptional repressors. Nat. Chem. Biol. 11, 207–213 (2015).

43. M. K. Sayeg, et al., Rationally Designed MicroRNA-Based Genetic Classifiers Target Specific Neurons in the Brain. ACS Synth. Biol. 4, 788–795 (2015).

44. H. Huang, et al., Oncolytic adenovirus programmed by synthetic gene circuit for cancer immunotherapy. Nat. Commun. 10, 4801 (2019).

45. D. Ma, S. Peng, Z. Xie, Integration and exchange of split dCas9 domains for transcriptional controls in mammalian cells. Nat. Commun. 7, 13056 (2016).

46. D. T. Gillespie, The chemical Langevin equation. J. Chem. Phys. 113, 297–306 (2000).

47. Y. Zhao, X. Shen, T. Tang, C.-I. Wu, Weak Regulation of Many Targets Is Cumulatively Powerful—An Evolutionary Perspective on microRNA Functionality. Mol. Biol. Evol. 34, 3041–3046 (2017).

48. Y. Zhao, et al., Regulation of Large Number of Weak Targets—New Insights from Twin-microRNAs. Genome Biol. Evol. 10, 1255–1264 (2018).

49. C. E. Vejnar, M. Blum, E. M. Zdobnov, miRmap web: comprehensive microRNA target prediction online. Nucleic Acids Res. 41, W165–W168 (2013).

50. V. Agarwal, G. W. Bell, J.-W. Nam, D. P. Bartel, Predicting effective microRNA target sites in mammalian mRNAs. eLife 4 (2015).

51. M. Kertesz, N. Iovino, U. Unnerstall, U. Gaul, E. Segal, The role of site accessibility in microRNA target recognition. Nat. Genet. 39, 1278–1284 (2007).

52. R. Denzler, et al., Impact of MicroRNA Levels, Target-Site Complementarity, and Cooperativity on Competing Endogenous RNA-Regulated Gene Expression. Mol. Cell 64, 565–579 (2016).

53. J. J. Gam, J. Babb, R. Weiss, A mixed antagonistic/synergistic miRNA repression model enables accurate predictions of multi-input miRNA sensor activity. Nat. Commun. 9 (2018).

54. K. F. Murphy, R. M. Adams, X. Wang, G. Balázsi, J. J. Collins, Tuning and controlling gene expression noise in synthetic gene networks. Nucleic Acids Res. 38, 2712–2726 (2010).

55. R. D. Dar, N. N. Hosmane, M. R. Arkin, R. F. Siliciano, L. S. Weinberger, Screening for noise in gene expression identifies drug synergies. Science 344, 1392–1396 (2014).

56. A. Aranda-Díaz, K. Mace, I. Zuleta, P. Harrigan, H. El-Samad, Robust Synthetic Circuits for Two-Dimensional Control of Gene Expression in Yeast. ACS Synth. Biol. 6, 545–554 (2017).

57. A. L. Szymczak, et al., Correction of multi-gene deficiency in vivo using a single “self-cleaving” 2A peptide–based retroviral vector. Nat. Biotechnol. 22, 589–594 (2004).

