## Supplemental Figures and Tables for "Characterizing microRNA-mediated modulation of gene expression noise and its effect on synthetic gene circuits"

### Supplementary Figures

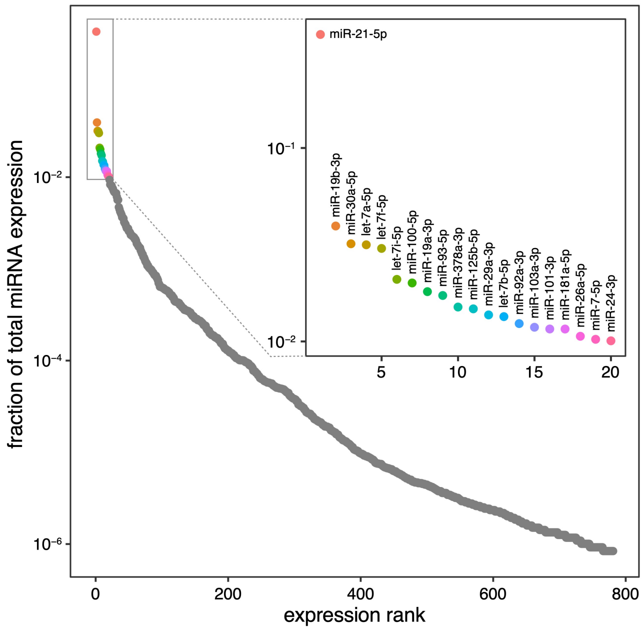


Fig. S1 miRNA expression profiles of HeLa cells measured by miRNA-seq. miRNAs that were involved in this study were labeled in color.

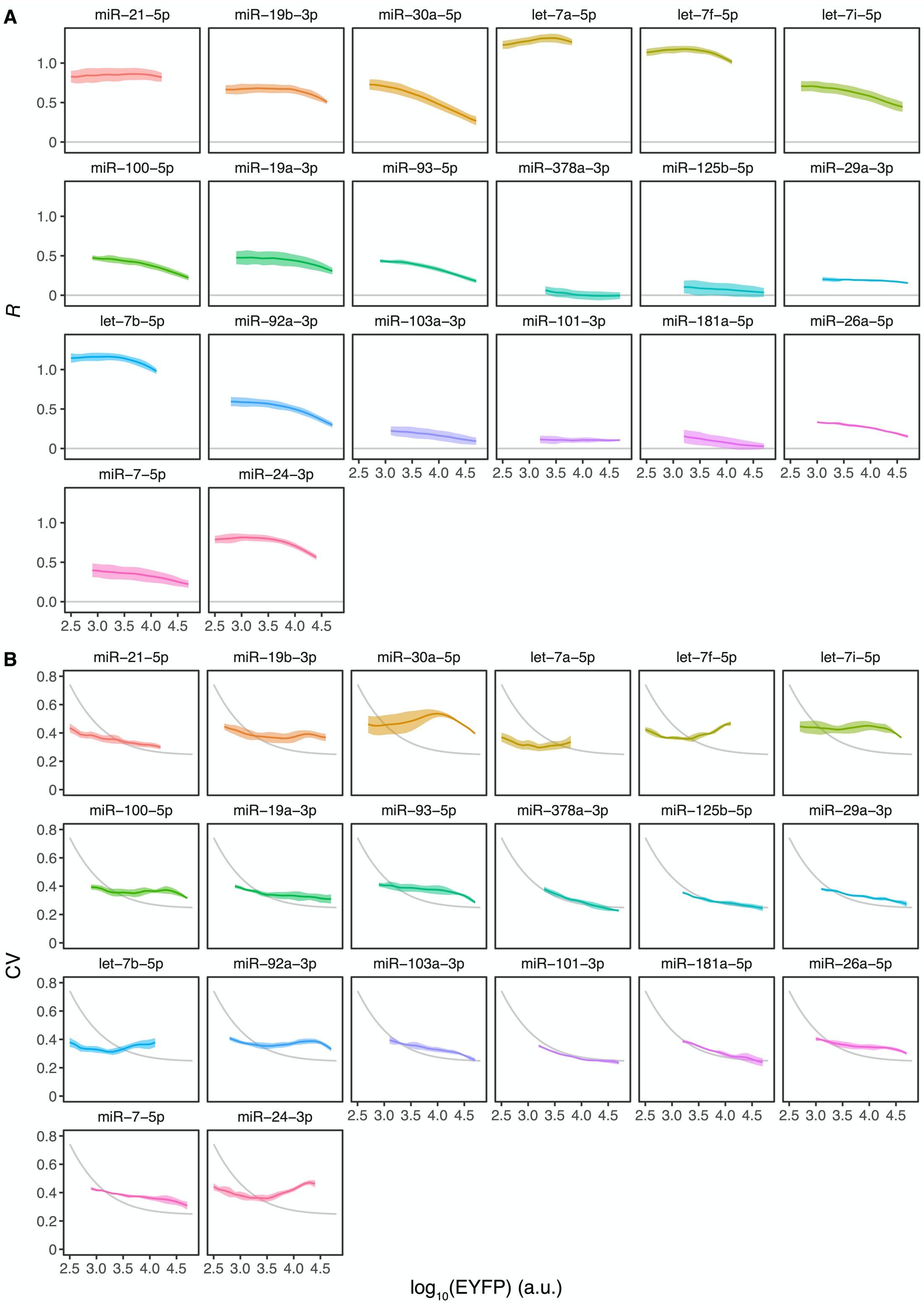


Fig. S2 (*A*) Repression strength and (*B*) noise level of dual reporters with a single MRE. Lines and shading show the mean ± SD of three independent replicates. Gray lines represent fitted results of reporters without MREs.

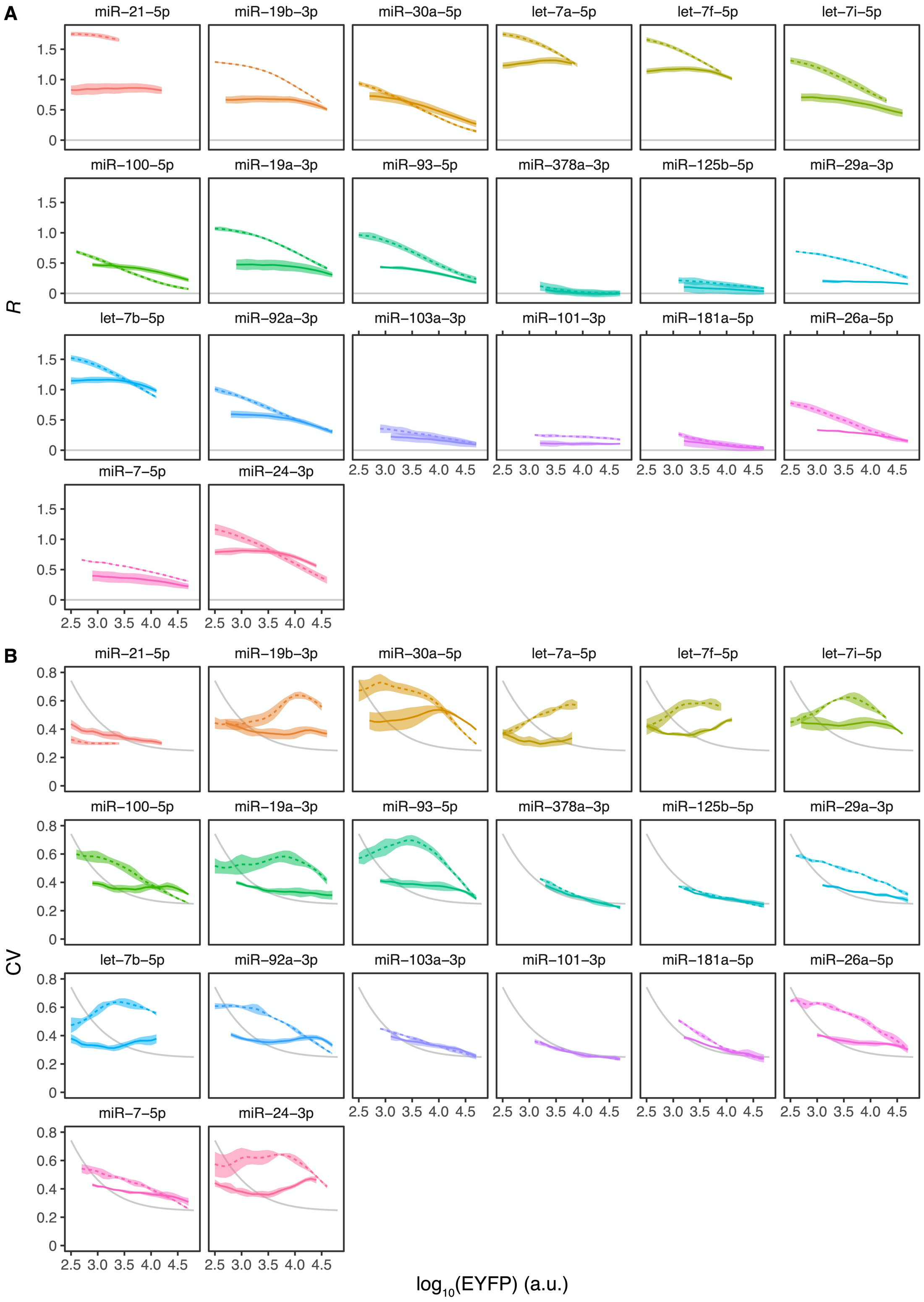


Fig. S3 (*A*) Repression strength and (*B*) noise level of dual reporters with triple MREs (colored dashed lines) in comparison with those with a single MRE (colored solid lines). Lines and shading show the mean ± SD of three independent replicates. Gray lines represent fitted results of reporters without MREs.

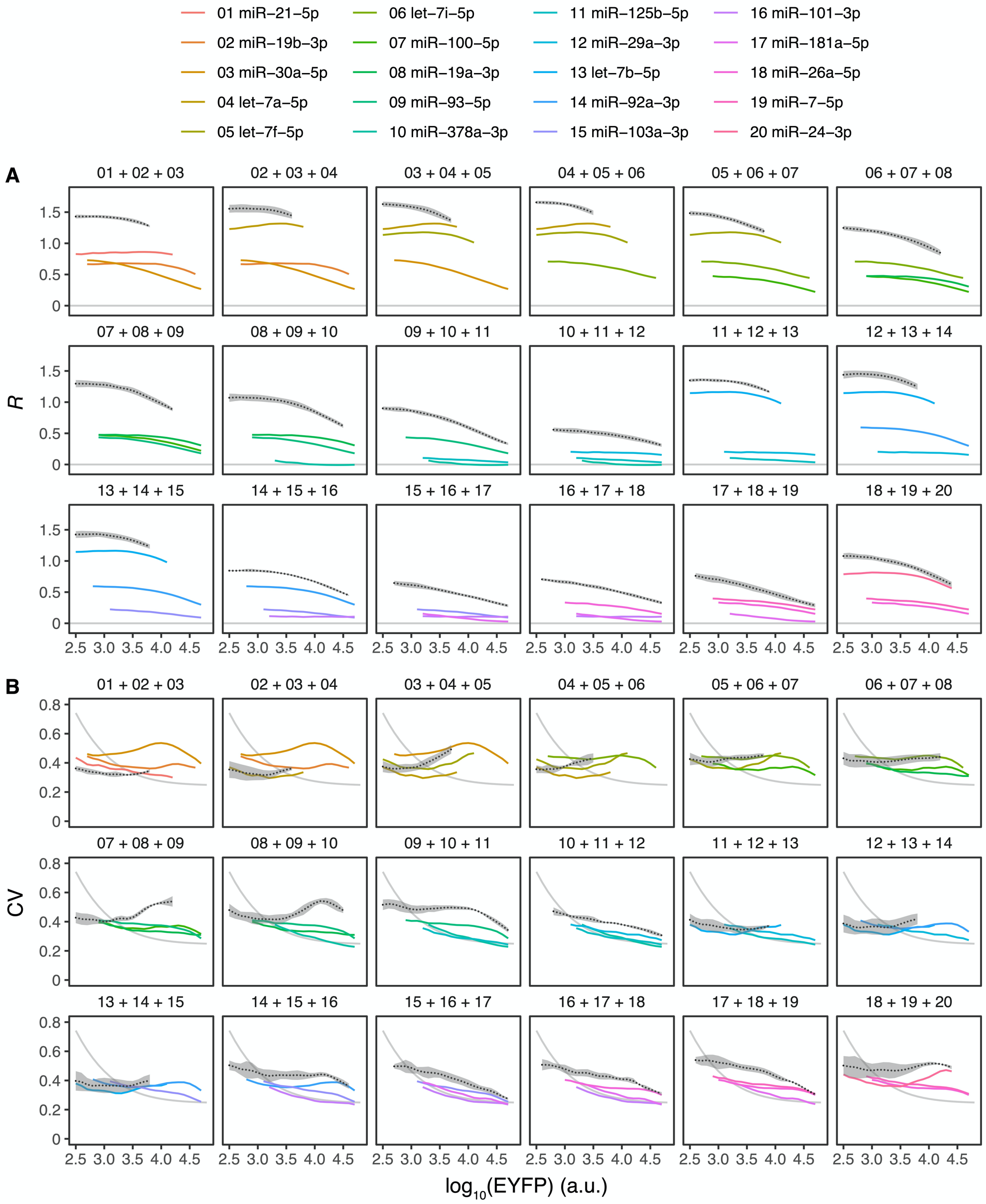


Fig. S4 (*A*) Repression strength and (*B*) noise level of dual reporters with composite MREs (black dotted lines) in comparison with those with a single MRE (colored solid lines, standard deviations not shown). Lines and shading show the mean ± SD of three independent replicates. Gray lines represent fitted results of reporters without MREs.

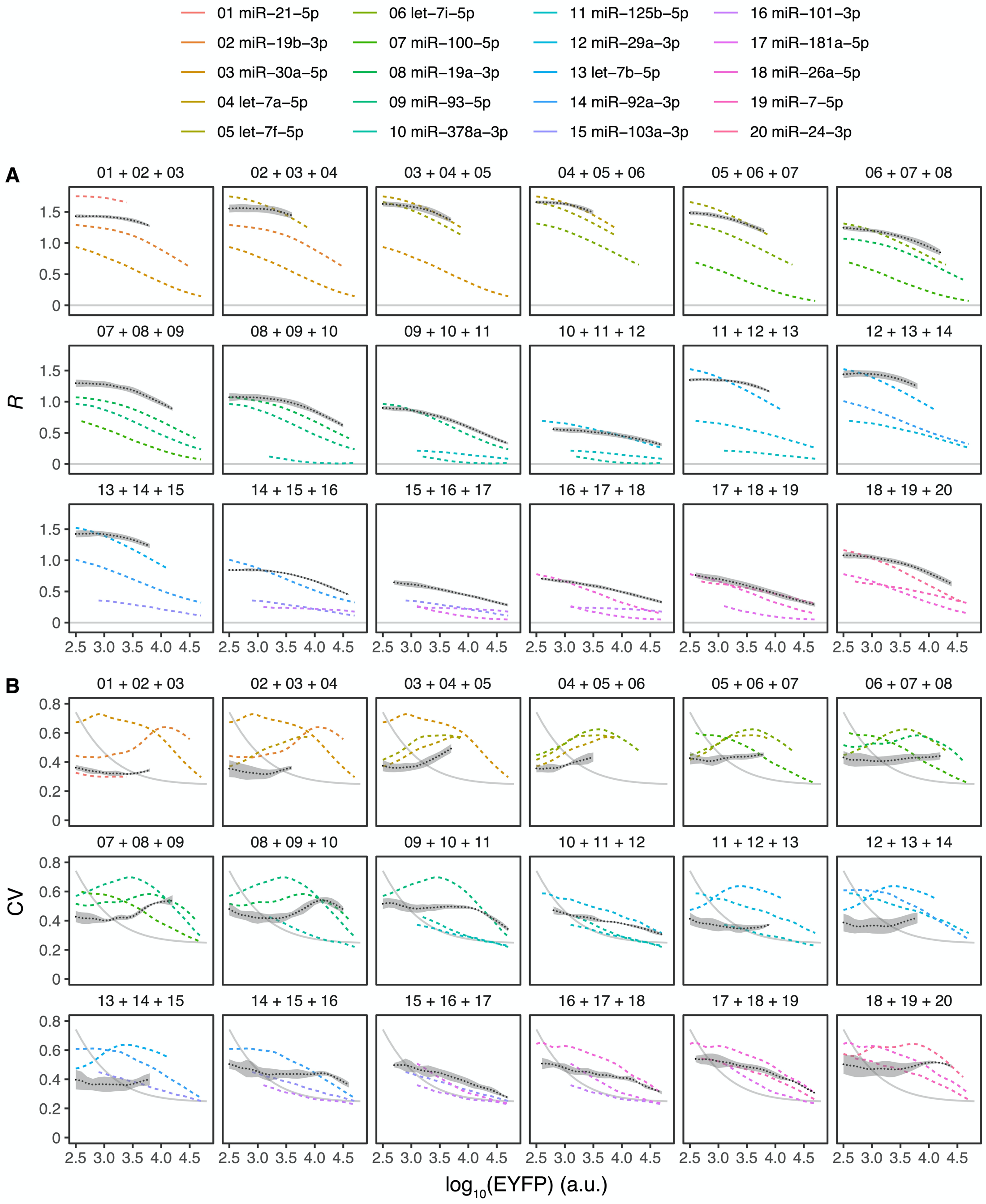


Fig. S5 (*A*) Repression strength and (*B*) noise level of dual reporters with composite MREs (black dotted lines) in comparison with those with triple MREs (colored dashed lines, standard deviations not shown). Lines and shading show the mean ± SD of three independent replicates. Gray lines represent fitted results of reporters without MREs.

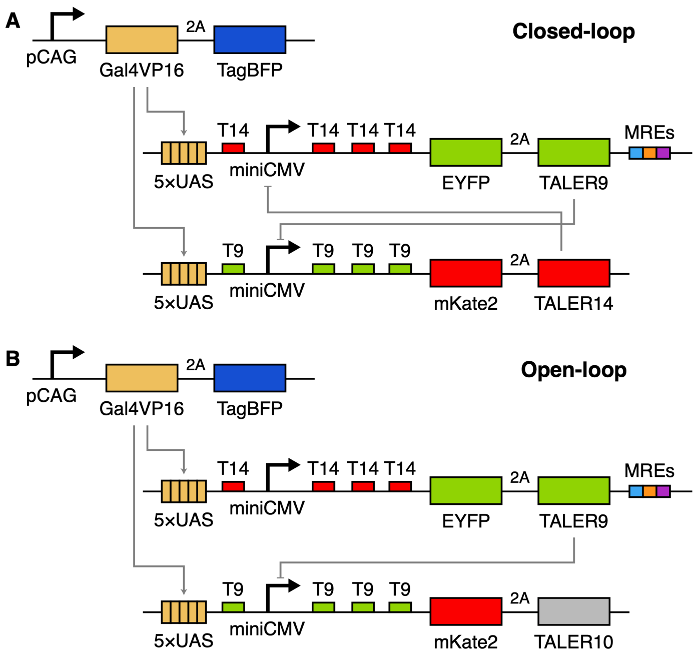


Fig. S6 Schematic diagrams of the (*A*) closed-loop TALER switches and (*B*) open-loop TALER circuits.

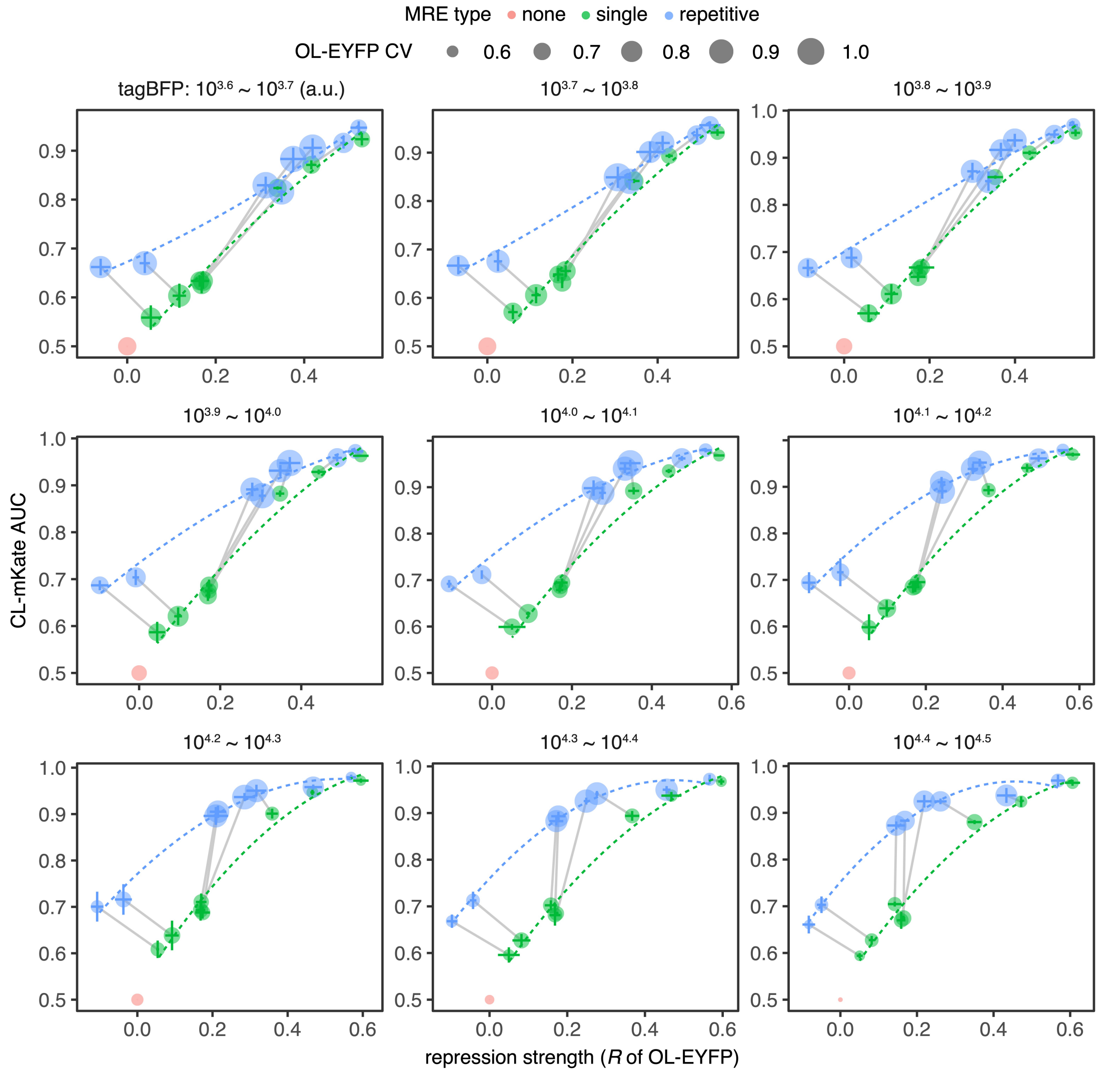


Fig. S7 The relationship between the repression strength of MREs in OL circuits and the AUCs classified by mKate2 in CL switches in different TagBFP bins. The size of the point represents the noise level (CV) of EYFP mediated by the MRE in OL circuits. Dashed curves represent the loess regression of the points. Each point shows mean ± SD with three independent replicates. Points that represent reporters with a single MRE or triple MREs of the same miRNA are connected by gray lines.

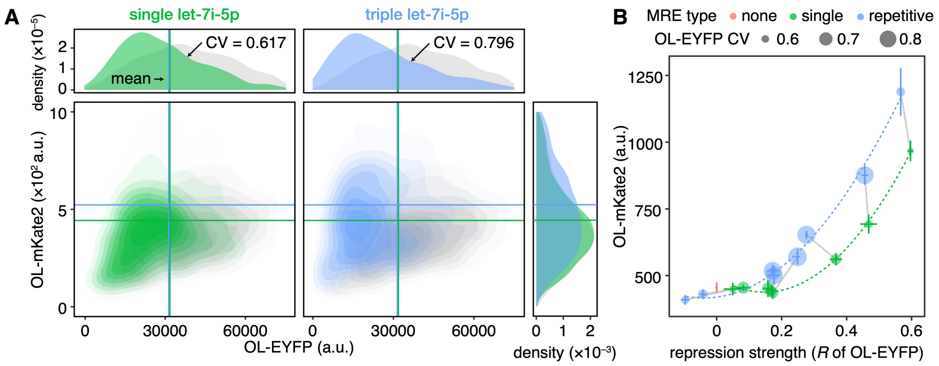


Fig. S8 (*A*) Joint and marginal distributions of EYFP and mKate2 in OL circuits with a single MRE or triple MREs of let-7i-5p. Colored lines represent the mean expression value of fluorescent protein in corresponding switches. (*B*) The relationship between repression strength and mKate2 mean expression in OL switches. The size of the point represents the noise level (CV) of EYFP mediated by the MRE in OL circuits. Dashed curves represent the loess regression of the points. Each point shows mean ± SD with three independent replicates. Points that represent reporters with a single MRE or triple MREs of the same miRNA are connected by gray lines.

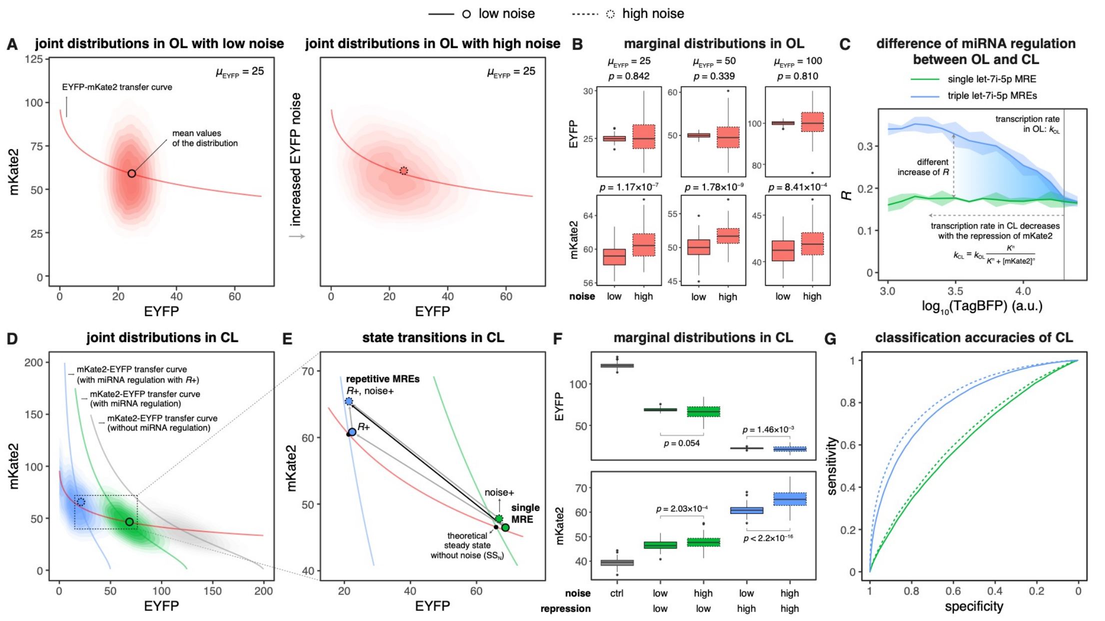


Fig. S9 Stochastic simulation results of the OL and CL circuits. (*A*) Joint distributions of EYFP and mKate2 in OL circuits with the mean value of EYFP ($\mu_{\mathrm{EYFP}}$) equals to 25. (*B*) Distributions of the mean values of EYFP and mKate2 in OL circuits with different $\mu_{\mathrm{EYFP}}$. (*C*) Repression strengths of the single and triple let-7i-5p MREs measured in OL circuits under different transcription rates (represented by the intensity of TagBFP). Lines and shading show the mean ± SD of three independent replicates. (*D*) Joint distributions of EYFP and mKate2 in CL switches with different miRNA regulatory conditions (single MRE: low noise and low repression strength, repetitive MREs: high noise and high repression strength). (*E*) State transition from single MRE to repetitive MREs. The mean values of EYFP and mKate2 in simulated CL switches with different miRNA regulatory conditions. *R*+ indicates the increase of the repression strength of miRNA and noise+ indicates the increase of the noise of EYFP. (*F*) Distributions of the mean values of EYFP and mKate2 in CL switches with different miRNA regulatory conditions. (*G*) ROC curves obtained when classifying cells by mKate2 between the condition with and without the regulation of miRNA in CL switches. In *A*, *D* and *E*, red curves represent the transfer curves from EYFP to mKate2; gray curves represent the transfer curves from mKate2 to EYFP without miRNA regulation; green curves represent the transfer curves from mKate2 to EYFP with miRNA regulation with low repression strength; blue curves represent the transfer curves from mKate2 to EYFP with miRNA regulation with high repression strength. All transfer curves are calculated by the deterministic equations. The cross points of the transfer curves (black points in *E*) represent the theoretical steady state without noise ($SS_N$). The colored dots represent the mean values of EYFP and mKate2 of the corresponding distribution. Mann–Whitney *U* tests of the mean EYFP or mKate2 expression levels were performed between different conditions in *B* and *F*, each condition with 100 samples. The significance levels were denoted as *p*.

As shown in *A*, the expression level of mKate2 is around the transfter curve from EYFP to mKate2 (red lines). The extent of mKate2 expression elevation is much higher than the that of mKate2 expression reduction due to the nonlinearity of the EYFP-mKate2 transfer curve. The expression of EYFP disperses more with the increase of EYFP noise, causing a more severe elevation of mKate2 at the low expression level of EYFP, which further leads to the increase of mKate2 mean values in OL circuits (*B*). The results are consistent across different $\mu_{\mathrm{EYFP}}$ values (*B*).

As shown in *C*, the transcription rate of EYFP mRNA is much lower in CL switches in comparison with the corresponding OL circuits because of the presence of mKate repression. Repetitive MREs usually exhibit stronger saturation effects in comparison with the single MRE (Fig. 3E and *C*), so in CL switches, where the repression of mKate2 on EYFP exists, the repression strength of the single MRE increases moderately in comparison with OL circuits, but that of the repetitive MREs increases remarkably due to the desaturation of miRNA. Thus, the transfer curve from mKate2 to EYFP alters from the green line to the blue line in *D*, leading to the transition of $SS_N$ (black points in *E*) to a lower-EYFP and higher-mKate2 state (*E*). Furthermore, we found that CL switches with low EYFP noise exhibited a state (solid points in *E*) quite near $SS_N$. However, in CL switches with high EYFP noise, the expression of mKate2 increases as shown in *A*, and it in turn represses the expression of EYFP, which brings about a state with lower EYFP and higher mKate2 (dashed points in *E*) deviating from $SS_N$. With the presence of both the increase of noise and repression strength as the real repetititve MREs behave, the alteration of $SS_N$ contributed by higher repression and the deviation from $SS_N$ contributed by higher noise synergistically promote the cell state transition to a lower-EYFP and higher-mKate state (*D*), which further leads to the difference of EYFP and mKate expression (*F*) and the improvement of classification accruracies (*G*). The transitions of mean values of the distribution under different conditions are shown in *E*.

### Supplementary Tables

#### Table S1. miRNAs and the sequences of the corresponding single MRE.

| Expression rank | Name | MRE sequence |
|---|---|---|
| 1 | miR-21-5p | CAACATCAGTCTGATAAGCT |
| 2 | miR-19b-3p | TCAGTTTTGCATGGATTTGCACA |
| 3 | miR-30a-5p | AGCTTCCAGTCGAGGATGTTTACA |
| 4 | let-7a-5p | AACTATACAACCTACTACCTCA |
| 5 | let-7f-5p | AACTATACAATCTACTACCTCA |
| 6 | let-7i-5p | TAACAGCACAAACTACTACCTCAA |
| 7 | miR-100-5p | CACAAGTTCGGATCTACGGGTT |
| 8 | miR-19a-3p | TCAGTTTTGCATAGATTTGCACA |
| 9 | miR-93-5p | CTACCTGCACGAACAGCACTTTG |
| 10 | miR-378a-3p | GCCTTCTGACTCCAAGTCCAGT |
| 11 | miR-125b-5p | TCACAAGTTAGGGTCTCAGGGA |
| 12 | miR-29a-3p | TAACCGATTTCAGATGGTGCTA |
| 13 | let-7b-5p | AACCACACAACCTACTACCTCA |
| 14 | miR-92a-3p | ACAGGCCGGGACAAGTGCAATA |
| 15 | miR-103a-3p | TCATAGCCCTGTACAATGCTGCT |
| 16 | miR-101-3p | TTCAGTTATCACAGTACTGTA |
| 17 | miR-181a-5p | ACTCACCGACAGCGTTGAATGTT |
| 18 | miR-26a-5p | AGCCTATCCTGGATTACTTGAA |
| 19 | miR-7-5p | AACAACAAAATCACTAGTCTTCCA |
| 20 | miR-24-3p | CTGTTCCTGCTGAACTGAGCCA |

#### Table S2. Parameter settings of flow cytometry experiments.

| Fluorescent protein | Laser (nm) | Filter | Photomultiplier tube voltage | |
|---|---|---|---|---|
| | | | For dual reporter | For TALER switch |
| TagBFP | 405 | 450/50 | 280 | 240 |
| mKate2 | 561 | 610/20 | 340 | 400 |
| EYFP | 488 | 530/30 | 210 | 285 |

#### Table S3. Parameters for simulations of Fig. 2B.

| Parameter | Red line | Green line | Blue line |
|---|---|---|---|
| $k_R$ | $5\times10^{-3}$ | $1.5\times10^{-2}$ | $1.5\times10^{-2}$ |
| $g_R$ | $1\times10^{-4}$ | | |
| $k_{T1}$ | $1\times10^{-3}$ | | |
| $k_{T2}$ | 0 | $2\times10^{-2}$ | $2.25\times10^{-2}$ |
| $g_{T1}$ and $g_{T2}$ | $1\times10^{-5}$ | | |
| $k_{1+}$ and $k_{2+}$ | $1\times10^{-4}$ | | |
| $k_{1-}$ | $5\times10^{-5}$ | | |
| $k_{2-}$ | / | $5\times10^{-5}$ | $5\times10^{-3}$ |
| $g_1$ and $g_2$ | $8\times10^{-5}$ | | |
| $\alpha_1$ | 1 | | |
| $\alpha_2$ | 0.5 | | |
| $k_{P1}$ | $1\times10^{-2}$ | | |
| $g_{P1}$ | $5\times10^{-6}$ | | |

#### Table S4. Reactions and parameters for stochastic simulations of TALER circuits.

| Parameter | | Value |
|---|---|---|
| $k_{TK}$ | | 0.1 |
| $g_{TK}$ | | 0.01 |
| $K_E$ and $K_K$ | | 50 |
| $m$ | | 0.5 |
| $n$ | | 1.56 |
| $k_{TE0}$ | | 1 |
| $g_{TE0}$ | | 0.05 |
| $R$ | $\mu_{\text{EYFP}} = 25$ in OL | 8 |
| | $\mu_{\text{EYFP}} = 50$ in OL<br>high repression in CL | 4 |
| | $\mu_{\text{EYFP}} = 100$ in OL | 2 |
| | low repression in CL | 1.6 |
| $t$ | low noise | 1 |
| | high noise | 0.01 |
| $k_{PE}$ and $k_{PK}$ | | 0.1 |
| $g_{PE}$ and $g_{PK}$ | | 0.01 |
